# How comparable across management goals are grassland monitoring methods?

**DOI:** 10.64898/2026.05.18.726054

**Authors:** Hunter Messick, Elinor M. Lichtenberg

## Abstract

**Questions:** Ecological monitoring, repeated collection of ecological data, is essential to document how ecosystems respond to change. In grasslands, different vegetation monitoring protocols are used across disciplines, making it difficult to address multiple management objectives or research questions. We asked four questions about how three common vegetation monitoring protocols compare. (1) How do the protocols differ in how they collect data? (2) How do the protocols differ in their utility? (3) In what ways do vegetation measurements quantitatively differ across protocols? (4) What are each protocol’s strengths?

**Location:** This study was conducted on working ranches in the Southern Great Plains with vegetation consisting mainly of native forbs and grasses.

**Methods:** We implemented three protocols at each site: (1) the Rangeland Analysis Platform (RAP), (2) the Grassland Effectiveness Monitoring (GEM) protocol, and (3) a typical pollinator ecology survey protocol. We qualitatively compared each protocol’s utility and quantitatively compared cover measurements that each produced.

**Results:** All three protocols displayed positive associations within cover categories, but differed in actual cover measurements. The RAP protocol, which uses remote sensing, measured the highest total vegetation cover. The GEM protocol, a line-point intercept method, had more capability to capture fine-scale cover patterns. The GEM protocol measured the most bare ground while the Pollinator protocol measured more forb coverage.

**Conclusion:** Fine-scale methods like the GEM protocol are most appropriate to address objectives that require capturing small patterns that would otherwise be overlooked with methods like quadrats or remote sensing. Remote sensing is advantageous when monitoring large areas or inaccessible land, but may over-estimate cover. The Pollinator protocol is best equipped to address questions regarding flower abundance and richness. Similarities among protocols can facilitate synergy across disciplines for more effective monitoring. We emphasize the importance of denoting a clear scale and scope of monitoring objectives before selecting methods.

## INTRODUCTION

Grasslands make up approximately 40% of the Earth’s terrestrial surface (Petermann & Buzhdygan, 2021). These systems serve as biodiversity hubs, particularly at temperate latitudes (Habel et al., 2013), and account for upwards of one third of terrestrial productivity (Vitousek, 2015). Beyond providing essential habitat for many species, grasslands serve as essential environmental regulators. Grasslands have immense carbon sequestration capacity and regulate erosion and water cycling and quality (Bai & Cotrufo, 2022; Bengtsson et al., 2019). Humans rely on grassland ecosystems for agriculture, particularly for livestock grazing (Liu, Hu, & Lu, 2024). Like many other biomes, grasslands are facing unprecedented declines in biodiversity and ecosystem services due to climate and land use change (Hooper et al., 2012; Liu, Hu, & Lu, 2024). In the Midwest US, for example, existing grasslands have been converted to corn and soy bean crops (Zhang et al., 2021). The importance of grasslands economically, environmentally, and culturally highlights the need to understand how these systems respond to human impacts. Ecological monitoring, long-term data collection with repeated measurements (Elzinga, Salzer, & Willoughby, 1999), has become imperative to record changes over time and inform management decisions. Monitoring can indicate responses to environmental conditions, provide early warning signs of ecosystem declines, improve understanding of ecosystem dynamics, and inform management decisions (Sparrow et al., 2020).

Grassland conservation and monitoring is a target for multiple disciplines and scientists studying various taxa, processes, and research questions. With a range of stakeholders and management goals comes similar variety in monitoring practices. Monitoring for plant conservation often focuses on documenting imperiled native plant species (Lavery et al., 2021). Ranchers and rangeland scientists record grassland canopy cover and heterogeneity to monitor livestock forage (Ge et al., 2025; Woods & Ruyle, 2015). Grassland bird populations rely on vegetation composition, structural heterogeneity, and patchiness (Hovick, Elmore, & Fuhlendorf, 2014; Hu et al., 2025). Pollination ecologists collect data on forb diversity, nesting resources provided by vegetation and ground cover, and pollinator communities (Collins et al., 2025; Zeng et al., 2023). It is also common to monitor erosion by describing ground cover based on organic litter or bare soil (Briske, 2017).

Monitoring has many benefits across disciplines; however, implementation comes with several challenges. First, monitoring is time-consuming and can be expensive. Some restoration projects overlook or underfund monitoring due to lack of resources or budgeting (Bernhardt et al., 2005). Efforts are often restricted to small spatial and temporal scales (Pringle et al., 2025). Monitoring often requires specialized skills in species identification or survey methods (Kallimanis et al., 2012). Furthermore, monitoring methods are often not standardized across disciplines, making it difficult to compare trends and address multiple management objectives across the greater landscape (Briske, 2017; Godínez-Alvarez et al., 2009). These challenges must be mitigated to facilitate efficient collection of ecological data.

We provide a formal methods comparison to offer insight into the applications and effectiveness of different monitoring approaches in grasslands. We compared three protocols that examine grassland plant community composition and structure: (1) the US Department of Agriculture’s (USDA) Rangeland Analysis remote sensing Platform (RAP), (2) US Fish and Wildlife Service’s (USFWS) Grassland Effectiveness Monitoring (GEM) protocol, and (3) a typical pollination ecology protocol. The RAP is an online database of spatial layers derived from satellite imagery of vegetation cover and above-ground biomass at 10-m and 30-m pixel resolutions (Allred et al., 2025). This platform aims to provide an accessible way for landowners, land managers, researchers, and conservationists to access vegetation and biomass data in US rangelands. The GEM protocol is an on-the-ground, line-point intercept protocol developed through a collaboration of federal and state agencies, non-profits, and private companies to effectively monitor grassland responses to practices like prescribed fires, grazing, herbicide application, plantings, and brush management (Matthews et al., 2023). This protocol offers a standardized monitoring procedure and aims to be accessible to and consistently applied by different organizations, agencies, and conservation programs. It seeks to overcome the historical inconsistencies of informal monitoring on private ranches (Woods & Ruyle, 2015). A typical vegetation survey used to study pollinator ecology and conservation, hereafter the “Pollinator protocol”, consists of on-the-ground, quadrat-based methods to record key aspects of grassland related to food and nesting resources for pollinators including canopy cover, ground cover, and floral richness and abundance (Giovanetti et al., 2021). We used one example, the methodology outlined in Collins et al. (2025).

Implementing monitoring depends on practitioners’ research questions, skills, resources, capacity, and study system. Taking these considerations into account, we answered four questions that may guide future practitioners in selecting an appropriate monitoring protocol for their objectives. (1) How do the protocols differ in how they collect data? We focused our comparison on whether data came from on-the-ground collection or remote sensing, and the resolution at which vegetation was measured. (2) How do the protocols differ in their utility? (3) In what ways do vegetation measurements quantitatively differ across protocols? Measurements assessed included functional vegetation cover, ground cover, bare ground, and flowering plant species richness. (4) What are each protocol’s strengths? To answer these questions, we implemented the RAP, GEM, and Pollinator protocols at the same sites. We compared resulting measurements of grassland canopy and ground cover both qualitatively and quantitatively. While we did not conduct a cost-benefit analysis, we documented total time required to conduct each survey to provide information on overall effort required for on-the-ground monitoring.

## METHODS

### Study system

This study was conducted on 19 working ranches in North Texas that are spread across Denton, Cooke, and Wise counties. Texas contains over 47 million hectares (117 million acres) of non-federal grazing lands, accounting for nearly 20% of grazing lands in the US (United States Department of Agriculture, 2020). At each site, we implemented all three protocols to allow for direct comparison. All sites were located within the Cross Timber ecoregion, which is characterized by a mosaic of tallgrass prairies and woodlands dispersed across rolling hills (Griffith et al., 2007). This ecoregion has highly variable precipitation, typically 686 - 965 mm per year. There are two distinct wet seasons, in the spring and fall. Summer temperatures range from 21 to 36° C. All sites were actively grazed by livestock. Sites were spatially independent, separated by at least 1.5 km from all other sites. Each site consisted of a relatively open area containing and surrounded by grassland vegetation. We marked the center of each site to ensure that the GEM and Pollinator transects originated from the same spot.

### Rangeland Analysis Platform (RAP)

The Rangeland Analysis Platform web application is an open-access spatial layer that contains vegetation and ground cover estimates obtained from satellite imagery (Fig. 1a) (Allred et al., 2025; https://rangelands.app/rap/). We used the 10-m cover layer and the cover types outlined in Table 1. We recorded the GPS coordinates of each site’s center (Garmin eTrex 22x) and used ArcGIS Pro to calculate the proportion of total vegetation, bare ground, and canopy gaps within a 100 m radius of each site’s center.

**Table 1.** Categories from the Rangeland Analysis Platform used to classify canopy and ground cover. A star (*) indicates that the category was excluded from analysis due to no or consistently very low coverage.

| **Canopy Cover** | **Ground Cover** |
| --- | --- |
| Perennial forb & grass | Bare ground |
| Annual forb & grass |  |
| Shrub* |  |
| Tree* |  |

**Figure 1.**
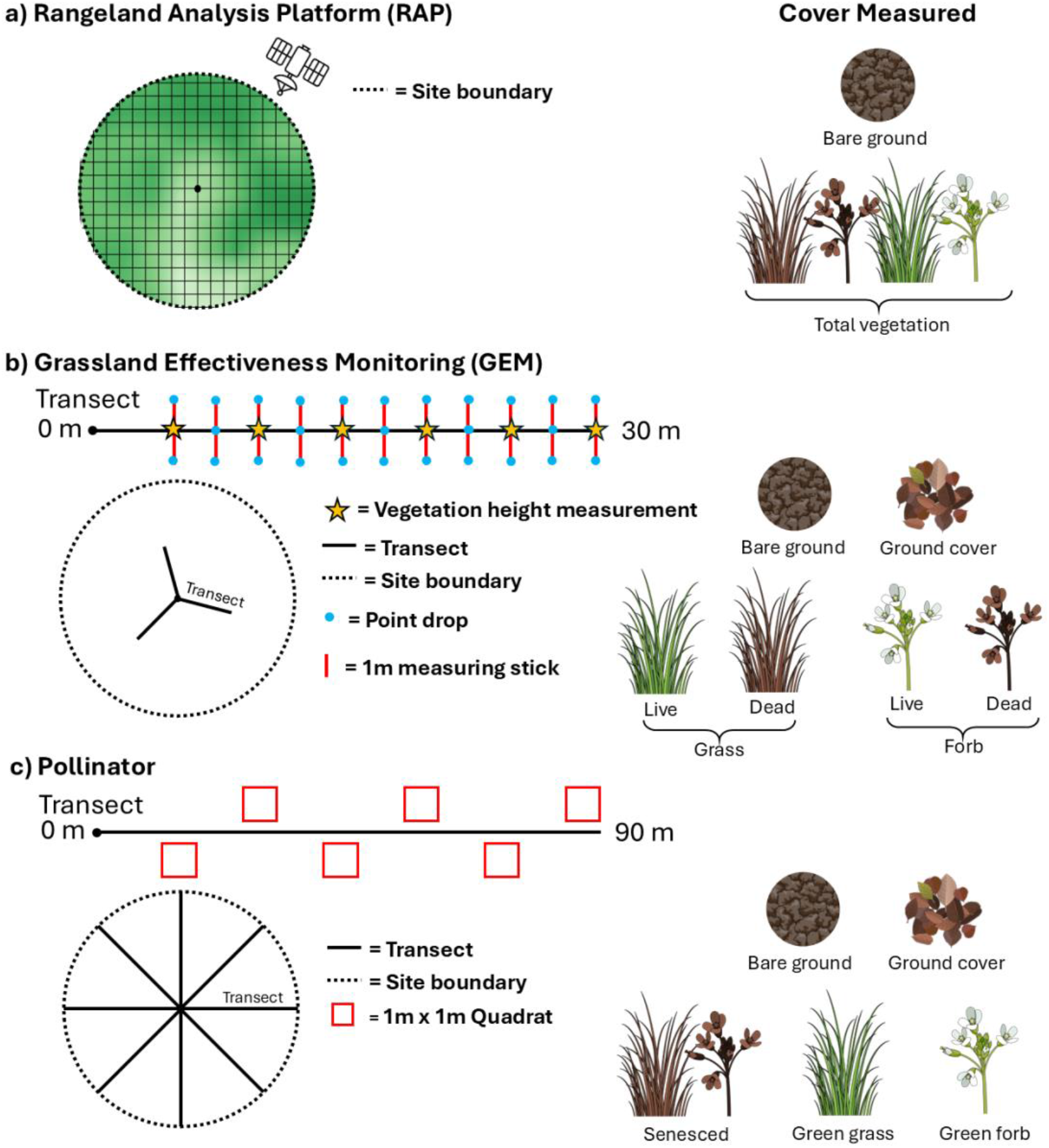
Schematic of the (a) Rangeland Analysis Platform, (b) Grassland Effectiveness Monitoring protocol, and (c) Pollinator protocol. Schematics are roughly to scale within but not across panels. Site boundaries refer to area sampled and not physical demarcations. Each icon on the right represents a cover type measured by the corresponding protocol. Icons sourced from ScienceFigures.org and used under Open Design License 1.0.

### Grassland Effectiveness Monitoring (GEM) protocol

The GEM protocol has three distinct “tiers,” with more complicated tiers requiring more detailed plant species identification (Matthews et al., 2023). We used the simplest tier, Tier 3, which is most accessible to non-scientists and researchers who do not have plant identification expertise (Fig. 1b). At each site, we selected a random starting azimuth then conducted three, 30 m step-point transects originating at the site’s center. The direction of first transect used the random azimuth and we subsequently added 120° and 240° from the original azimuth for the second and third transects. Starting five meters from the site center (to avoid trampled vegetation), we placed a meter stick centered on and perpendicular to the transect. We dropped a pin at the left side, center, and right sides of the meter stick and recorded vegetation interceptions from the uppermost canopy to the ground. We categorized vegetation interceptions into functional groups (Table 2) and recorded whether the intercepted vegetation was alive or senesced. We also categorized ground cover. If the soil beneath the ground cover layer had embedded material, we recorded this as a separate “ground” layer. If there was no embedded material beneath the ground cover layer, the ground layer was denoted as “soil”. We used the interceptions to calculate proportional cover by each category. At each end of the meter stick, we measured litter depth with a ruler. We also measured the largest herbaceous and woody canopy gaps that intercepted the meter stick. Starting from the right-most side of the meter stick, we categorized herbaceous canopy gaps as no gap (gap < 25 cm long), 25-49 cm, 50 - 99 cm, 100 - 199 cm, ≥ 200 cm. We estimated woody canopy gaps as not present (i.e., no woody plant encountered), < 5 m, 5 - 25 m, > 25 m. We repeated these measurements every 2.5 m along the 30 m transect totaling 12 locations per transect and 36 per site. We recorded woody and herbaceous plant height every five meters along the transect. At the center pin, we measured to the nearest centimeter the heights of the tallest herbaceous plant and the tallest woody plant within 30.5 cm of the pin. If woody plants were taller than two meters, we visually approximated to the nearest 30 cm, and plants with a height greater than or equal to 18 m were recorded as 18 meters tall. We also recorded whether woody plants were living or senesced.

**Table 2.** Categories used in the Grassland Effectiveness Monitoring (GEM) protocol to classify canopy, overlying ground, and ground-level cover. A star (*) indicates that the category was excluded from analysis due to no or consistently very low coverage.

| **Canopy Cover** | **Ground Cover** | **Ground (Including Embedded)** |
| --- | --- | --- |
| Forb | Herbaceous litter | Soil |
| Grass | Loose woody litter* | Gravel* |
| Sedge* | Vagrant lichen* | Cobble* |
| Vine* | Deposited soil/rock fragment* | Stone* |
| Shrub* | Other litter* | Boulder* |
| Tree* | No interception | Bedrock* |
| Cactus* |  | Basal plant base* |
| Yucca* |  | Lichen* |
| Succulent* |  | Moss* |
| No interception |  | Cyanobacteria* |
|  |  | Duff* |
|  |  | Embedded litter* |
|  |  | Trash* |
|  |  | Water* |

To make the GEM protocol more comparable to the Pollinator protocol, we opted to identify forbs to species if the pin intercepted a flower. This is not included in the original GEM protocol. Typically, the GEM protocol tier we implemented only identifies plants to functional group, but practitioners may identify to species if the plant is known.

### Pollinator protocol

Following the methods used by the Lichtenberg Lab at the University of North Texas (Collins et al., 2025) (Fig. 1c), we measured vegetation cover, ground cover, and flowers in 1 × 1 m quadrats along eight, 90 m transects radiating in the cardinal and ordinal directions. For each transect, we placed the quadrat every 15 m totaling six quadrats per transect and 48 quadrats per site. To facilitate cover estimation, we divided the quadrat into 25 squares with string. We visually estimated each category’s cover (Table 3) to the nearest half square, and used these counts to estimate proportional cover by each category. Some of the Pollinator protocol’s ground cover categories correspond to elements of the GEM protocol’s ground layer. We also identified flowering plant species in each quadrat and counted floral abundance for each species. All species were photo-vouchered and uploaded to an iNaturalist project (Lichtenberg, 2021).

**Table 3.** Categories used in the Pollinator protocol to classify canopy and ground cover. A star (*) indicates that the category was excluded from analysis due to no or consistently very low coverage.

| **Canopy Cover** | **Ground Cover** |
| --- | --- |
| No canopy | Bare ground |
| Dead/senesced vegetation | Moss* |
| Green forb | Prostrate green vegetation* |
| Green grass | Litter |
| Standing wood* | Manure* |
| Tree canopy* | Stems* |
|  | Rock* |
|  | Wood* |

### Data analysis

Multiple vegetation and ground cover categories were never or rarely encountered, all woody canopy gaps were > 25 m, and we encountered a woody plant taller than two meters only five times, so these were not further considered (Tables 1-3). To determine how protocols differed in canopy and ground cover estimates, we implemented generalized linear mixed models with the glmmTMB package (Brooks et al., 2017) in R (R Core Team, 2025) and used an α of 0.05. Each model included cover as a proportion with protocol as a fixed effect and site as a random effect to account for repeated sampling. We used logistic regressions weighted by the number of observations used to calculate proportional cover under the given protocol. We developed a standardized approach to allow for comparisons across each method because the three protocols have different purposes and thus measure similar but different vegetation and ground cover. We measured total vegetation proportional cover by combining annual and perennial forbs and grasses under the RAP protocol, living and dead forbs and grasses under the GEM protocol, and senesced vegetation, green forbs, and green grasses under the Pollinator protocol. We also compared bare ground, which was a single variable under the RAP and Pollinator protocols, and combined no ground cover and bare ground for the GEM protocol. We additionally compared four finer-resolution proportional cover metrics between the GEM and Pollinator protocols: live/green forbs, live/green grass, no canopy, and (herbaceous) litter. The GEM protocol divides the canopy into vertical layers; the above analyses combined interceptions from both overstory and understory.

The overstory provides information about dominant vegetation or dominant functional groups at a site. The GEM protocol explicitly measures this layer, while the Pollinator protocol’s ocular approach combines overstory with partial views of the understory. To determine how cover of each functional group (forbs and grasses) compared between the GEM protocol’s overstory estimates and the Pollinator protocol’s estimates, we used generalized linear mixed models similar to those described above.

The RAP and GEM protocols measure canopy gaps, while the Pollinator protocol does not. Before regressing each herbaceous canopy gap category on protocol (as described above), correcting for multiple comparisons following Holm (1979), we implemented a simulation to estimate canopy gap size categories from the Pollinator protocol’s no cover proportion. We generated a series of quadrats randomly populated with TRUE or FALSE values for “no canopy” (TRUE = no canopy, FALSE = canopy present), using the uniform distribution and dividing each quadrat into 5 × 5 cm squares. From each random quadrat, we randomly selected a row and categorized the largest number of consecutive TRUE values into a GEM gap category. Next, we calculated the proportion of TRUE values for each quadrat to estimate the Pollinator protocol’s “no canopy” cover. To determine if the proportion of no canopy could be used to predict canopy gap size, we built random forest models using the ranger package (Wright, Wager, & Probst, 2026). We fit each model with a sample of quadrats equally distributed across the canopy gap categories. We fit the first model with a sample of five quadrats (one per gap category), and we increased the sample in 5-quadrat increments before stopping at 100 quadrats (20 per gap category). These quadrat counts span the range used by projects that measure vegetation to study pollinator biodiversity or conservation (Collins et al., 2025; Higginson & Dover, 2021). We used three metrics to assess model performance. (1) Accuracy, the percentage of correct predictions, compares actual versus predicted values for the data points used to train the model. (2) Cramer’s V measures the strength of the relationship between two categorical variables (here, observed and predicted gap category). (3) Out-of-bag error uses bootstrapping to measure the model’s accuracy with data points that were not used to train the model. Out-of-bag error indicates a model’s predictive performance.

## RESULTS

### How do the protocols differ in how they collect data and their utility?

All three protocols were able to measure plant functional group composition but varied in the information they provided. Only the Pollinator protocol identified plants to species, although it focused on flowering forbs. An alternative version of the GEM protocol does identify all plants to species but requires additional expertise. Both the GEM and Pollinator protocols classify plants into broad functional groups (e.g., grass, forb, shrub, tree). The RAP protocol does not distinguish between grasses and forbs, but instead between perennial and annual herbaceous plants.

The GEM and RAP protocols measured grassland structural heterogeneity via canopy gap measurements, which the Pollinator protocol does not include. The GEM protocol measured vegetation height and litter depth, while the other two protocols did not.

The protocols differed in the resolution they capture data at. At the finest scale, the GEM protocol relies on physical intersections with a pin during line-point intercept surveys. The Pollinator protocol relies on ocular estimates in quadrats, so each measurement reflects cover at the scale of centimeters to meters depending on quadrat size. Because we counted by half square, this was 200 cm^2^. The RAP protocol uses satellite imagery to measure cover in 10 × 10 m pixels, losing a significant amount of fine-scale variation in the landscape.

For a full survey, and accounting for site-level variation, the GEM protocol had a mean survey time of 45 minutes while the Pollinator protocol had a mean survey time of 132 minutes. This averaged 15 minutes per transect for the GEM protocol and 17 minutes per transect for the Pollinator protocol, with high variation in the latter depending on how many flowers were in the quadrats. The RAP protocol does not require any fieldwork but does require familiarity with spatial analysis. We used the website to extract cover data for each site, which was done in one day. Faster methods are available using ArcGIS Pro or other spatial analysis platforms, which do require some expertise. Once the data were downloaded, calculating proportion cover was straightforward and quick.

### Do vegetation measurements differ among protocols?

Vegetation measurements differed among protocols (Appendix S1, S2). Total canopy cover was highest with the RAP protocol and lowest with the Pollinator protocol (Fig. 2a). In contrast, the GEM protocol measured the most bare ground while the RAP protocol indicated the least (Fig. 2b). The Pollinator protocol and GEM protocol did not differ in green canopy cover, but the Pollinator protocol recorded more forb cover than the GEM protocol (Fig. 3). The Pollinator protocol also recorded less senesced vegetation than the GEM protocol (Fig. 3). The decreased senesced cover was driven largely by green forbs (Fig. 3). On the ground, the Pollinator protocol measured higher litter cover than the GEM protocol (Fig. 4).

**Figure 2.**
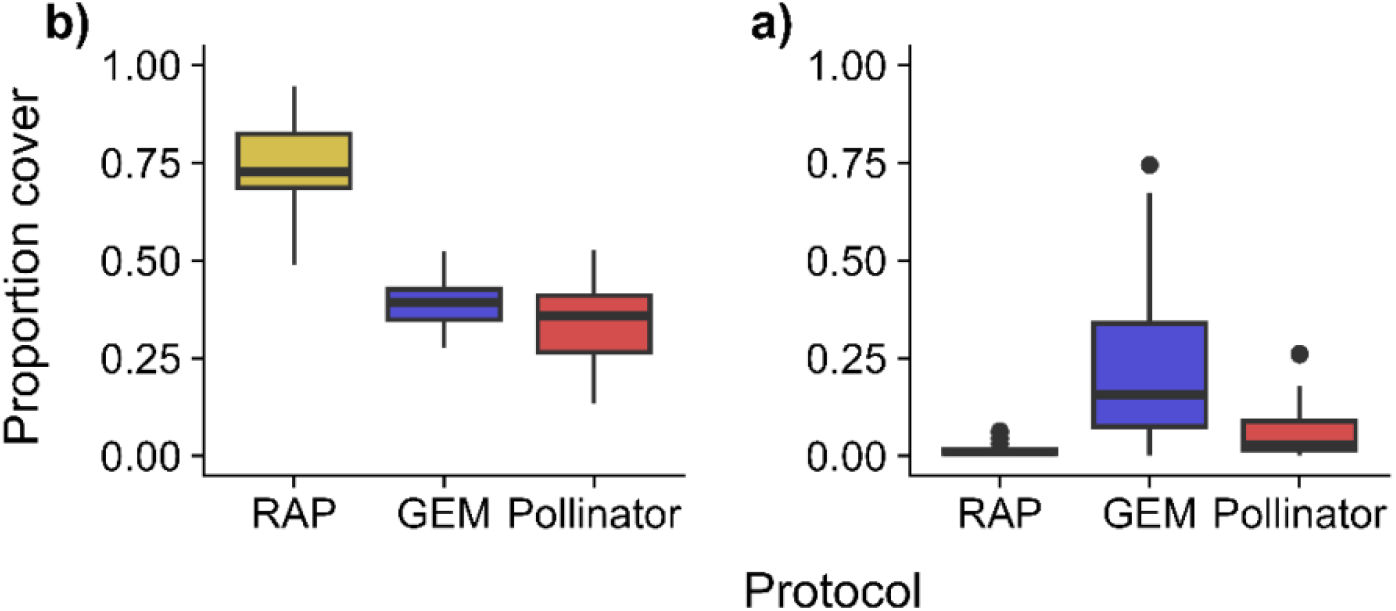
a) Total vegetation and b) bare ground measured by the RAP, GEM, and Pollinator protocols.

**Figure 3.**
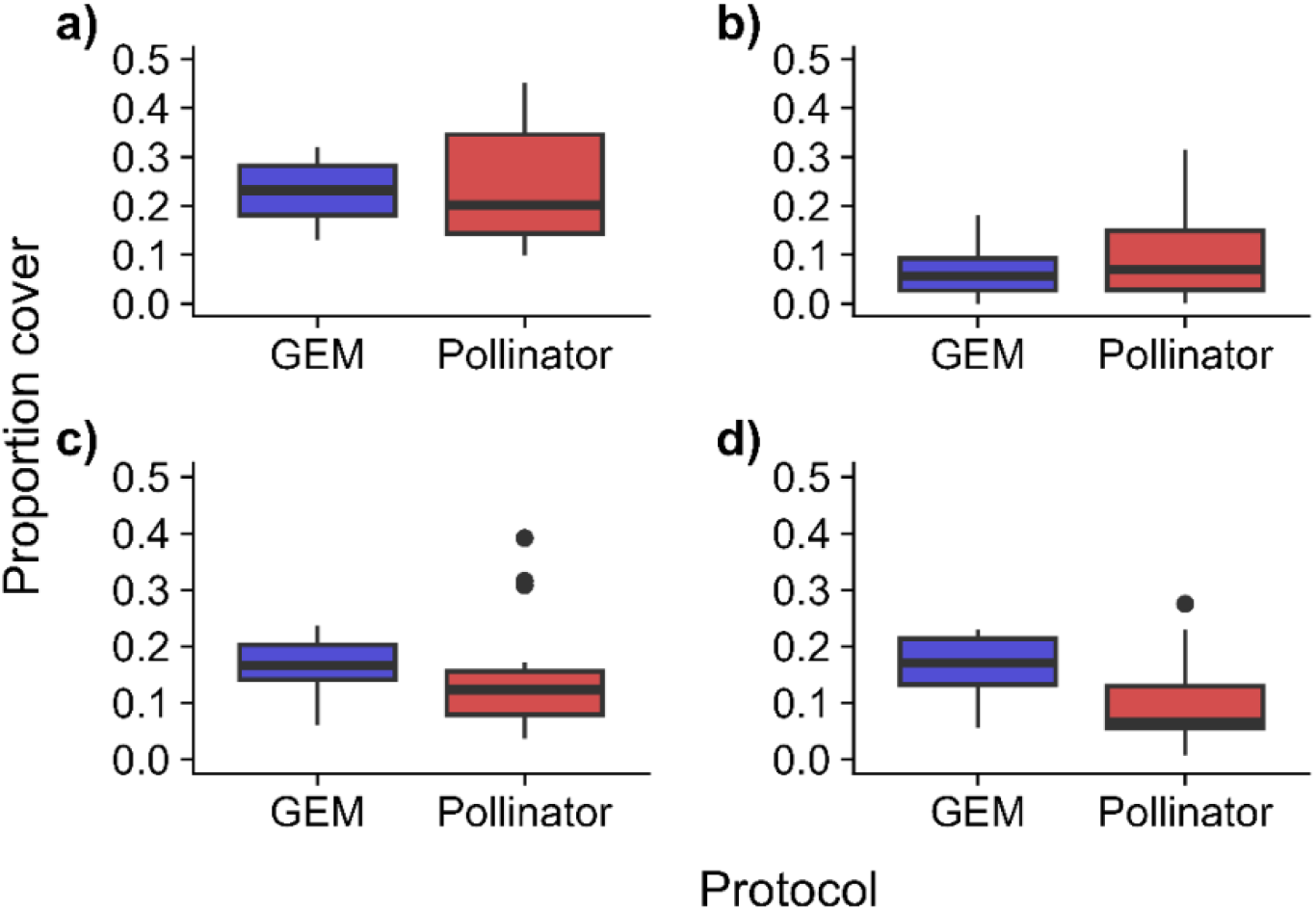
a) Green canopy, b) green forb, c) green grass, and d) senesced vegetation measured by the GEM and Pollinator protocols.

**Figure 4.**
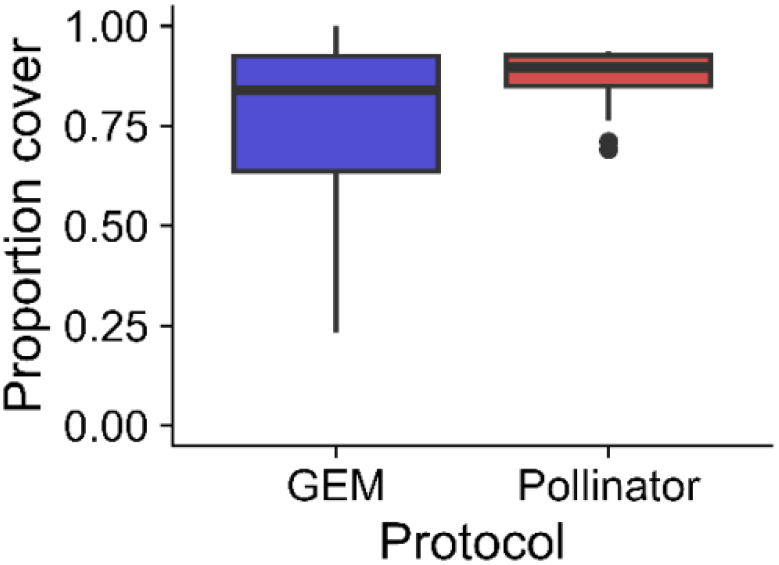
Litter cover measured by the GEM and Pollinator protocols.

The GEM protocol is designed to estimate relative cover of each plant functional group using just overstory intercepts. The GEM protocol overstory measurements were dominated by grass while the Pollinator protocol measured more forbs across the entire canopy (Appendix S3, S4).

Our random forest model indicated that we cannot reliably predict canopy gap categories from proportion of no canopy in Pollinator quadrats. The model was fairly accurate with training data, with accuracy always above 75% (Appendix S5a) and Cramer’s V of approximately 0.75 (Appendix S5b). As the number of quadrats included in the model increased, accuracy decreased and Cramer’s V stayed relatively constant. However, out-of-bag error was typically around 40%, indicating that the model correctly predicted only 60% of instances (Appendix S5c). With only a few quadrats, as is sometimes done with pollinator monitoring, prediction became incorrect most of the time. We thus compared canopy gap measurements only between the RAP and GEM protocols. The GEM protocol consistently recorded more small canopy gaps (<25 cm and 25 cm – 50 cm) and fewer medium gaps (50 cm – 100 cm gaps; Appendix S1, S6).

Although the protocols differed in the values measured for many cover types, they tended to capture similar overall patterns in vegetation cover and ground cover (Appendix S7). Bare ground estimates strongly positively correlated across all protocols (*r* range 0.75-0.90). Total vegetation cover moderately positively correlated between the GEM and RAP protocols (*r* = 0.65). Green canopy, senesced vegetation, and green grass moderately positively correlated between the GEM and Pollinator protocols (*r* = 0.53, 0.52, 0.44, respectively). Green forb estimates strongly positively correlated between the GEM and Pollinator protocols (*r* = 0.91).

The modified GEM protocol only counted 36 flowers from 11 plant species. The Pollinator protocol counted 6033 flowers from 80 species. All plant species recorded by the GEM protocol were also recorded by the Pollinator protocol.

## DISCUSSION

This study showed how monitoring protocols differ in how they collect data, their utility, and their quantitative measurements of cover. We also identified each protocol’s strengths. The GEM protocol focused more on functional group composition, while the Pollinator protocol emphasized flowering plants. The RAP protocol focused on total vegetation, but distinguished by life cycle (perennial versus annual) rather than functional group (grasses versus forbs). The three protocols captured similar patterns in canopy and ground cover but differed in how much canopy and ground cover they indicated was present. The remote-sensing-based RAP protocol measured noticeably more canopy cover than the two on-the-ground protocols. The Pollinator protocol recorded relatively more forb cover, while the GEM protocol recorded relatively more green grass. The GEM protocol was less time-intensive than the Pollinator protocol but covered a smaller area.

Methodologies often differ in their estimates of the same cover type (Briske, 2017). Our comparison of total vegetation suggests that remote sensing approaches such as the RAP protocol may over-estimate cover, and that such web-based products need further ground-truthing. Despite this current limitation, such products can be useful for tracking change over time. Previous research agrees with our finding that line-point intercept methods measure similar or higher canopy cover than ocular methods (here, quadrats) (Godínez-Alvarez et al., 2009). The line-point intercept approach relies on physical interceptions with a pin; therefore, it acts at a very fine spatial resolution. Ocular estimation acts at a lower resolution, directly impacting the magnitude of cover estimates. We sampled during summer and thus senesced vegetation was relatively uncommon and patchily distributed. The line-point intercept method was more likely to intercept individual stems of senesced vegetation that would otherwise be overlooked in a landscape dominated by green canopy. By slightly zooming out, quadrats likely reflected the broader pattern. It is crucial to consider the scale needed to address a specific research question. If practitioners aim to study small-scale patterns, using a line-point intercept method may be most appropriate. One such case is “microsites” or “micro patches” that are only centimeters in size. Soil microsites can have large effects on the plant community (Stover & Henry, 2018). Micro edges among microsites can serve as transition zones, providing niche spaces for microbial organisms and plants (Dann et al., 2018; Kowalski & Henry, 2020). On the other hand, if the researcher is interested in relative cover over a larger spatial area and does not need to capture small-scale variation or answer highly local questions, the RAP protocol is the best approach.

Both the GEM and Pollinator protocols distinguish between living and senesced vegetation, and the GEM protocol further divides senesced vegetation into functional groups. Examining a plant community’s senescence patterns can provide information concerning plant development and responses to environmental stress. Senescence is partially dictated by seasonal and hormonal cues as a plant ages, so tracking it allows practitioners to monitor plant development in response to environmental conditions or management (Jibran et al., 2017). Environmental stressors such as drought, pesticide use, or extreme temperatures can catalyze premature senescence (Kim et al., 2020; Munné-Bosch, 2025; Sedigheh et al., 2011). Land managers using adaptive management also monitor senescence of plant functional groups as indicators of livestock forage quality. Understanding senescence patterns additionally allows practitioners to examine plant community temporal dynamics and responses to treatments or environmental effects (Leong & Roderick, 2015). The GEM protocol is best suited to address monitoring or research questions about plant senescence because it records senescence at the greatest detail.

Recording vegetation height can provide valuable information about the resources provided by and structure of vegetation. Although livestock tend to prefer short, green grasses (Pauler et al., 2020), tall grasses can serve as a source of biomass in periods of drought and during the winter when preferred forage is limited (Briske, 2017). In this case, tall grasses buffer a drought’s effect on herbivore populations. Recording information like plant height can give practitioners an idea of the quality of resources present in the grassland. Vegetation height is additionally important for many grassland animal species, like birds. Access to perches can improve grassland regeneration by increasing seed rain and seedling establishment (Gan et al., 2025). Taller grassland vegetation also provides nest cover and increased resource availability for ground-nesting and cereal-associated bird species (Faria et al., 2024). The GEM protocol was the only protocol that measured plant height, but this can be easily added to other on-the-ground protocols like the Pollinator protocol.

The Pollinator protocol focuses on food and nesting resources for pollinators, and measures flowering plant species richness and flower abundance. Although we adapted the GEM protocol to note flower intercepts, it recorded only a fraction of the flowers that the Pollinator protocol found. Floral species richness can be informative beyond pollination ecology. Forb composition can directly impact livestock nutrition and grazing behavior. Livestock have grazing preferences that include certain forb species (Hesselmann et al., 2025), and pastures with greater native forb abundance can enhance cattle weight gain (Prigge et al., 2024). Although the GEM protocol we followed in this study (Tier 3) did not record forb species richness or abundance, other versions that require greater plant identification expertise do record intercepted plants, but not flowers, to species (Tiers 1 and 2). Note that forbs will bloom only at specific times of year and may not bloom every year so forb species richness may be a poor indicator of floral availability (de Vries et al., 2026).

A limitation of this study is that we cannot definitively state which protocol comes closest to recording “true” cover values. Measuring true values at the necessary spatial scale for monitoring habitat would be prohibitively time intensive. Across ecological research, it is uncommon to estimate aspects of a habitat with 100% accuracy. Similarly, scaling from local measurements to system-level patterns or processes is a well-recognized challenge (Chacón-Labella et al., 2023; Qiu & Cardinale, 2020). The different cover values we found with smaller-scale methods (the GEM and Pollinator protocols) versus a large-scale approach (the RAP protocol) further demonstrate this challenge. Ecological systems are complex and unique; as such, practitioners should prioritize using methods most appropriate for the scale and scope of their questions, as well as within their budget, capacity, and resources.

Lack of standardized monitoring methods across disciplines has hindered addressing multiple management objectives and efficiently collecting data. We showed that current methods for monitoring grasslands are not completely interchangeable. However, all provided a core set of data and a general indication of what the other protocols would record. All measured vegetation cover and bare ground, and measurements positively correlated across protocols. This means that values measured through one protocol can potentially be used as rough estimates for cover types that were otherwise not recorded in another protocol. Small adaptations can also provide greater utility outside a protocol’s main purpose. For example, the Pollinator protocol could use fewer transects or quadrats when time is limited, or the GEM protocol could space points further apart along a longer transect in highly heterogeneous vegetation or to cover a larger area. Measuring vegetation height or litter depth can be added to any on-the-ground grassland monitoring protocol with little extra effort. Combining on-the-ground sampling via the GEM or Pollinator protocol with remote sensing such as the RAP protocol can help overcome the latter’s lack of detail while covering a broader spatial area and reducing field time. For specific tasks, however, certain protocols are the best option. If a monitoring project involves a range of stakeholders that differ in expertise and management goals, we recommend the GEM protocol because it collects a variety of information while requiring limited expertise. This protocol can be made more complex or simple depending on users’ plant identification expertise (Matthews et al., 2023). If on-the-ground sampling is not possible due to limited resources, inaccessible sites, or large spatial extent, the RAP protocol will be most useful. Any use of RAP, however, should keep in mind its likely over-estimation of cover. Lastly, if flowers are of particular interest, we recommend using the Pollinator protocol.

Different protocols can also be used in conjunction with one another. For example, practitioners could identify potential sites using the RAP protocol, before performing on-the-ground surveys using the GEM or Pollinator protocol. Because each protocol presents unique utility and strengths, they each provide a range of benefits and limitations as well as varying time and effort requirements. While many collect similar data, specific objectives or finer scale questions may require a larger investment or a modification from standard protocols to provide details needed to answer those questions. Because these protocols vary in their strengths, we stress the importance of clearly articulating management goals and research questions prior to selecting monitoring protocols.

## Acknowledgements

We thank CP Cattle Company, the Dixon Water Foundation, Silver Spur Ranch, Todd and Stephanie Underwood, and Wilson Land and Cattle for ranch access. We thank Avery Pearson, Aidan McKinnis, Julian Moore, Jada Martinez, and Jodi Knight for assistance in data collection and plant identification. Lastly, we thank Don Wilhelm, Daniel Bunting, and the GEM program for their contributions and development of the GEM survey protocols that we tested in the field. This material is based upon work that is supported by the National Institute of Food and Agriculture, U.S. Department of Agriculture, under award number 2022-38640-37488 through the Southern Sustainable Agriculture Research and Education program under subaward number OS23-162. USDA is an equal opportunity employer and service provider.

## Supporting Information

Additional supporting information may be found online in the Supporting Information section:

Appendix S1. Coefficients and likelihood ratio test statistics for the models investigating the effect of protocol on canopy cover, ground cover, and canopy gap measurements.

Appendix S2. Test statistics for post-hoc Tukey tests of pairwise differences among protocols for models that compared all three protocols.

Appendix S3. Coefficients and likelihood ratio test statistics for the model comparing green forbs and green grass measured by the GEM protocol’s overstory hits with the Pollinator protocol.

Appendix S4. Canopy cover estimates recorded only in the overstory for the GEM protocol compared with the Pollinator protocol estimates.

Appendix S5. Random forest model accuracy, Cramer’s V, and out-of-bag error for models fit with 5 to 100 quadrats.

Appendix S6. Canopy gaps recorded by the GEM and RAP protocols.

## Notes

### Competing Interest Statement

The authors have declared no competing interest.

