## Appendix for "How comparable across management goals are grassland monitoring methods?"

**SUPPLEMENTAL MATERIAL FOR: Are grassland monitoring methods comparable across different management goals?**

**Appendix S1.** Coefficients and likelihood ratio test statistics for the models investigating the effect of protocol on canopy cover, ground cover, and canopy gap measurements. Canopy gap analysis corrected for multiple comparisons using the method described by Holm (1979).

| | **Model term** | **Coefficient** | **Coefficient standard error** | **$\chi^2$** | **df** | ***p*-value** |
|---|---|---|---|---|---|---|
| Canopy cover | **Model: Total vegetation ~ Protocol** | | | | | |
| | Intercept | -0.43 | 0.08 | | | |
| | Protocol: Pollinator | -0.27 | 0.07 | 1911.20 | 2 | **<0.0001** |
| | Protocol: RAP | 1.53 | 0.04 | | | |
| | **Model: Green canopy ~ Protocol** | | | | | |
| | Intercept | -1.23 | 0.08 | | | |
| | Protocol: Pollinator | 0.07 | 0.08 | 0.84 | 1 | 0.36 |
| | **Model: Green forb ~ Protocol** | | | | | |
| | Intercept | -3.11 | 0.27 | | | |
| | Protocol: Pollinator | 0.50 | 0.12 | 15.69 | 1 | **<0.0001** |
| | **Model: Green grass ~ Protocol** | | | | | |
| | Intercept | -1.64 | 0.08 | | | |
| | Protocol: Pollinator | -0.17 | 0.10 | 3.16 | 1 | 0.08 |
| | **Model: Senesced vegetation ~ Protocol** | | | | | |
| | Intercept | -1.66 | 0.09 | | | |
| | Protocol: Pollinator | -0.67 | 0.12 | 38.82 | 1 | **<0.0001** |
| Ground cover | **Model: Bare ground ~ Protocol** | | | | | |
| | Intercept | -1.56 | 0.33 | | | |
| | Protocol: Pollinator | -1.77 | 0.15 | 1038.7 | 2 | **<0.0001** |
| | Protocol: RAP | -3.41 | 0.13 | | | |
| | **Model: Litter ~ Protocol** | | | | | |
| | Intercept | 1.37 | 0.27 | | | |
| | Protocol: Pollinator | 1.01 | 0.12 | 75.198 | 1 | **<0.0001** |
| | **Model: <25 cm canopy gaps ~ Protocol** | | | | | |

| | | | | | | |
|---|---|---|---|---|---|---|
| Canopy gaps | Intercept | 3.05 | 0.51 | | | |
| | Protocol: RAP | 2.49 | 0.18 | 167.79 | 1 | **<0.0001** |
| | **Model: 25 – 50 cm canopy gaps ~ Protocol** | | | | | |
| | Intercept | -3.26 | 0.54 | | | |
| | Protocol: RAP | -3.67 | 0.27 | 213.47 | 1 | **<0.0001** |
| | **Model: 50 cm – 100 cm canopy gaps ~ Protocol** | | | | | |
| | Intercept | -4.82 | 0.56 | | | |
| | Protocol: RAP | -1.69 | 0.38 | 15.89 | 1 | **<0.0001** |
| | **Model: 100 cm – 200 cm canopy gaps ~ Protocol** | | | | | |
| | Intercept | -6.87 | 1.00 | | | |
| | Protocol: RAP | -0.16 | 0.75 | 0.05 | 1 | 0.83 |
| | **Model: > 200 cm canopy gaps ~ Protocol** | | | | | |
| | Intercept | -7.22 | 1.09 | | | |
| | Protocol: RAP | 1.02 | 1.02 | 1.40 | 1 | 0.24 |

**Appendix S2**. Test statistics for post-hoc Tukey tests of pairwise differences among protocols for models that compared all three protocols.

| Comparison | Z | *p*-value |
|---|---|---|
| **Model: Total vegetation ~ Protocol** | | |
| Pollinator vs. GEM | -3.70 | **<0.001** |
| RAP vs. GEM | 39.82 | **<0.001** |
| RAP vs. Pollinator | 25.08 | **<0.001** |
| **Model: Bare ground ~ Protocol** | | |
| Pollinator vs. GEM | -6.00 | **<0.001** |
| RAP vs. GEM | -5.50 | **<0.001** |
| RAP vs. Pollinator | -3.35 | **0.002** |

**Appendix S3.** Coefficients and likelihood ratio test statistics for the models comparing green forbs and green grass measured by the GEM protocol's overstory with the Pollinator protocol.

| **Model term** | **Coefficient** | **Coefficient standard error** | $\chi^2$ | **df** | ***p*-value** |
|---|---|---|---|---|---|
| **Model: Green forb ~ Protocol** | | | | | |
| Intercept | -2.19 | 0.31 | | | |
| Protocol: Pollinator | -0.56 | 0.13 | 18.61 | 1 | **<0.0001** |
| **Model: Green grass ~ Protocol** | | | | | |
| Intercept | -0.31 | 0.14 | | | |
| Protocol: Pollinator | -1.58 | 0.11 | 263.98 | 1 | **<0.0001** |

**Appendix S4.** Canopy cover estimates recorded only in the overstory for the GEM protocol compared with the Pollinator protocol estimates.

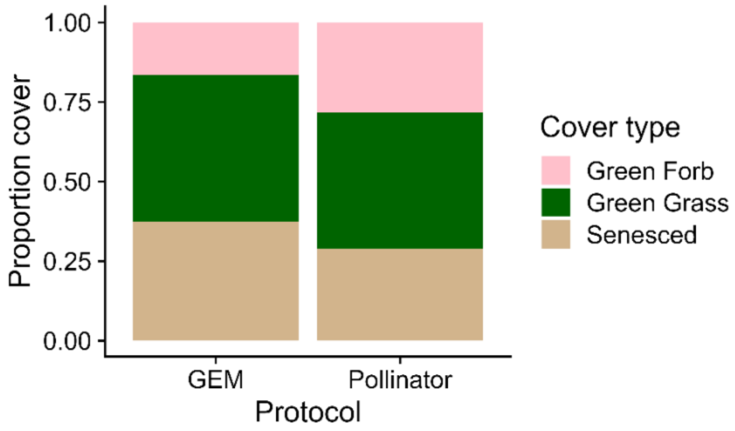

**Appendix S5.** Random forest model (a) accuracy, (b) Cramer's V, and (c) out-of-bag error for models fit with 5 to 100 quadrats. Each model contained the same number of simulated quadrats per canopy gap category.

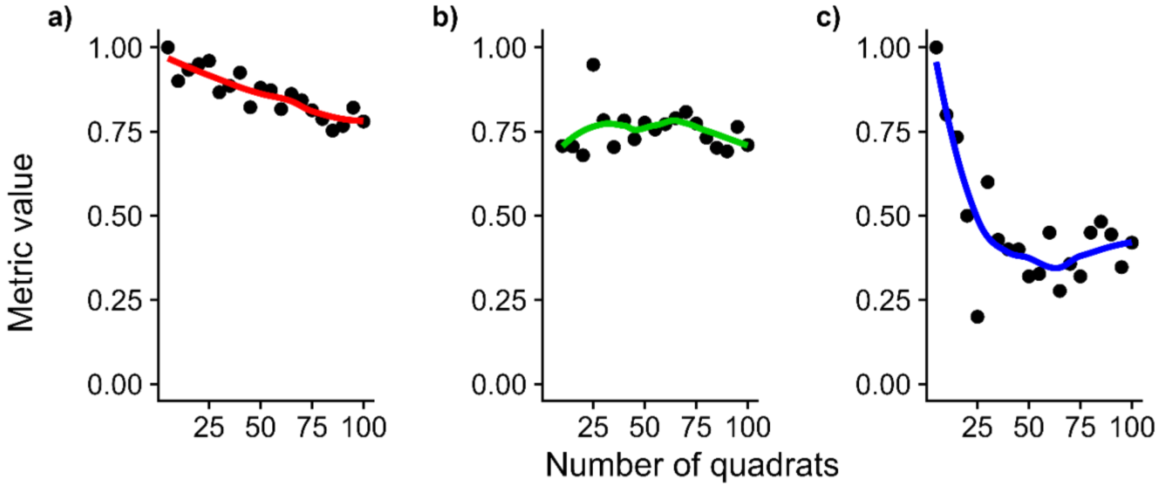

**Appendix S6.** Canopy gaps recorded by the GEM and RAP protocols.

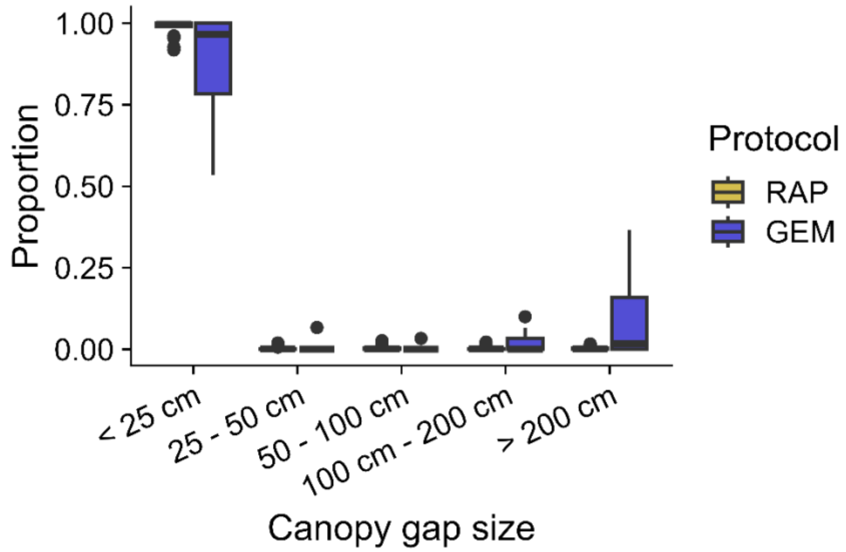

**Appendix S7.** Correlations among canopy and ground cover estimates within protocol pairs.

| **Cover type** | **Protocol pair** | | **S** | ***r*** | ***p*-value** |
|---|---|---|---|---|---|
| Total vegetation | GEM | Pollinator | 839.13 | 0.369 | 0.11 |
| | GEM | RAP | 461.67 | 0.653 | **0.002** |
| | Pollinator | RAP | 908.84 | 0.317 | 0.17 |
| Green canopy | GEM | Pollinator | 601.45 | 0.528 | **0.01** |
| Green forb | GEM | Pollinator | 126.64 | 0.905 | **<0.0001** |
| Green grass | GEM | Pollinator | 743.18 | 0.441 | 0.052 |
| Senesced vegetation | GEM | Pollinator | 639.44 | 0.519 | **0.02** |
| Bare ground | GEM | Pollinator | 134.75 | 0.899 | **<0.0001** |
| | GEM | RAP | 327.49 | 0.754 | **0.0001** |
| | Pollinator | RAP | 241.27 | 0.819 | **<0.0001** |
| Litter | GEM | Pollinator | 322.85 | 0.757 | **0.0001** |
